# Batch Correction via Cross-Modal Anchoring

**DOI:** 10.64898/2026.11.12.766280

**Authors:** Wassim Jaoui, Kjong-Van Lehmann

## Abstract

Batch correction is essential for integrating molecular cohorts, yet unimodal correction cannot reliably separate batch effects from biology without explicit covariates. Here we introduce anchor-guided batch correction, a label-free method that uses a comparatively stable paired omic layer to protect biology while correcting a batch-affected assay, requiring no predefined clinical covariates and returning full feature-level profiles. Across seven standard cohorts from The Cancer Genome Atlas, controlled confounding benchmarks, role-reversed transcriptomic workflows, and simulations, our method removed almost all platform variance in the standard cohorts, outperforming ComBat in biological preservation and CpG recovery under confounding. In the imbalanced breast cancer benchmark, where ComBat collapsed subtype retrieval (AUC 0.37) and attenuated differential methylation recovery (73.2%), anchor-guided correction preserved subtype structure (AUC 0.72) and recovered 82.8% of reference CpG signals. Running in three seconds on 1,091 tumours on a single CPU without external training data, cross-modal anchoring offers a fast, practical solution for paired multi-omic harmonization.

## Introduction

Paired multi-omic profiling captures coordinated biological programmes alongside variation specific to each molecular layer [1–3]. However, integrating multi-omic cohorts across laboratories, clinical recruitment waves, and evolving measurement technologies is frequently compromised by severe technical batch effects [4, 5]. Effective data harmonization must therefore satisfy two competing objectives: eliminating technical structure while preserving biological heterogeneity [6, 7]. Established unimodal correction methods, such as empirical-Bayes ComBat [8], estimate location and scale shifts from the affected assay. Control-gene approaches estimate unwanted variation from genes assumed to be biologically invariant [9]. In clinical cohorts, rare-subtype studies, and longitudinal platform transitions, patient composition can differ across technical groups [4, 5, 10]. When the relevant biology is unmeasured or fully confounded with batch, the affected assay alone cannot reliably distinguish technical offsets from biological divergence. Treating those biological differences as batch effects erases signal [6, 7, 10]. Supervised covariate protection requires measured biological covariates and cannot explicitly protect unannotated subtypes, plastic cell states, or unmeasured continuous gradients such as tumour purity and immune infiltration [8, 10].

Paired multi-omic designs offer a principled statistical solution to this identifiability problem. Existing methods learn joint latent factors for multi-omic integration [11, 12]. MultiBaC predicts missing assay profiles across studies [13]. Our method uses paired assays to protect biology during target correction. In practice, technical artefacts are frequently asymmetric across molecular layers: a cohort may undergo a major platform transition in one assay, such as a DNA methylation array upgrade or a sequencing workflow change, while its paired molecular counterpart remains comparatively stable. We designate the less distorted modality as the *anchor* and the batch-affected modality requiring harmonization as the *target*. Shared cross-modal structure can be learned from paired profiles through a regularized linear mapping, supplying a continuous, multivariate reference for biologically warranted inter-batch differences without predefined clinical or subtype covariates. Because the target modality also captures layer-specific biological programmes absent from the anchor, unpredicted target variation must not be equated with technical noise. The anchor should constrain alignment using shared biology, while target-specific variation is modified only when evidence supports a technical adjustment.

Here, we introduce anchor-guided batch correction, which uses paired measurements to protect biological differences while estimating technical adjustments in a low-dimensional space. These adjustments are applied to the original feature profiles, producing corrected data in the full feature space while preserving all variation outside the corrected subspace. Across 33 cancer types from The Cancer Genome Atlas (TCGA) [1, 14], we characterize technical asymmetry between paired RNA and methylation, then evaluate biological preservation in seven standard cohorts and controlled breast-cancer confounding benchmarks. Reversing the modality roles in glioblastoma demonstrates methylation anchoring of RNA correction, evaluated against housekeeping controls and ComBat. Simulations establish the operating limits and test preservation of both shared and modality-private biology.

## Results

### Paired RNA exposes methylation platform distortion

In TCGA, paired molecular profiles revealed a marked asymmetry between DNA methylation measured on HumanMethylation27 (HM27) and HumanMethylation450 (HM450) arrays and paired RNA expression. In breast cancer (BRCA), the arrays separated the DNA-methylation profiles of 1,091 tumours into distinct platform groups (**Figure 1a**), whereas the same patients overlapped more extensively in paired RNA expression (**Figure 1b**). An analysis of 33 TCGA cancer types identified ten projects profiled on both arrays (**Figure 1c, Supplementary Table 1**). Platform separation was 3.2-to 13.5-fold greater in methylation than in RNA, despite identical patient groupings in both assays **(see Methods)**. Colouring profiles by biological subtype revealed platform structure superimposed on subtype organization, illustrated here in breast cancer (**Figure 1d**), while subtype structure remained visible in paired RNA (**Figure 1e**).

**Figure 1.**
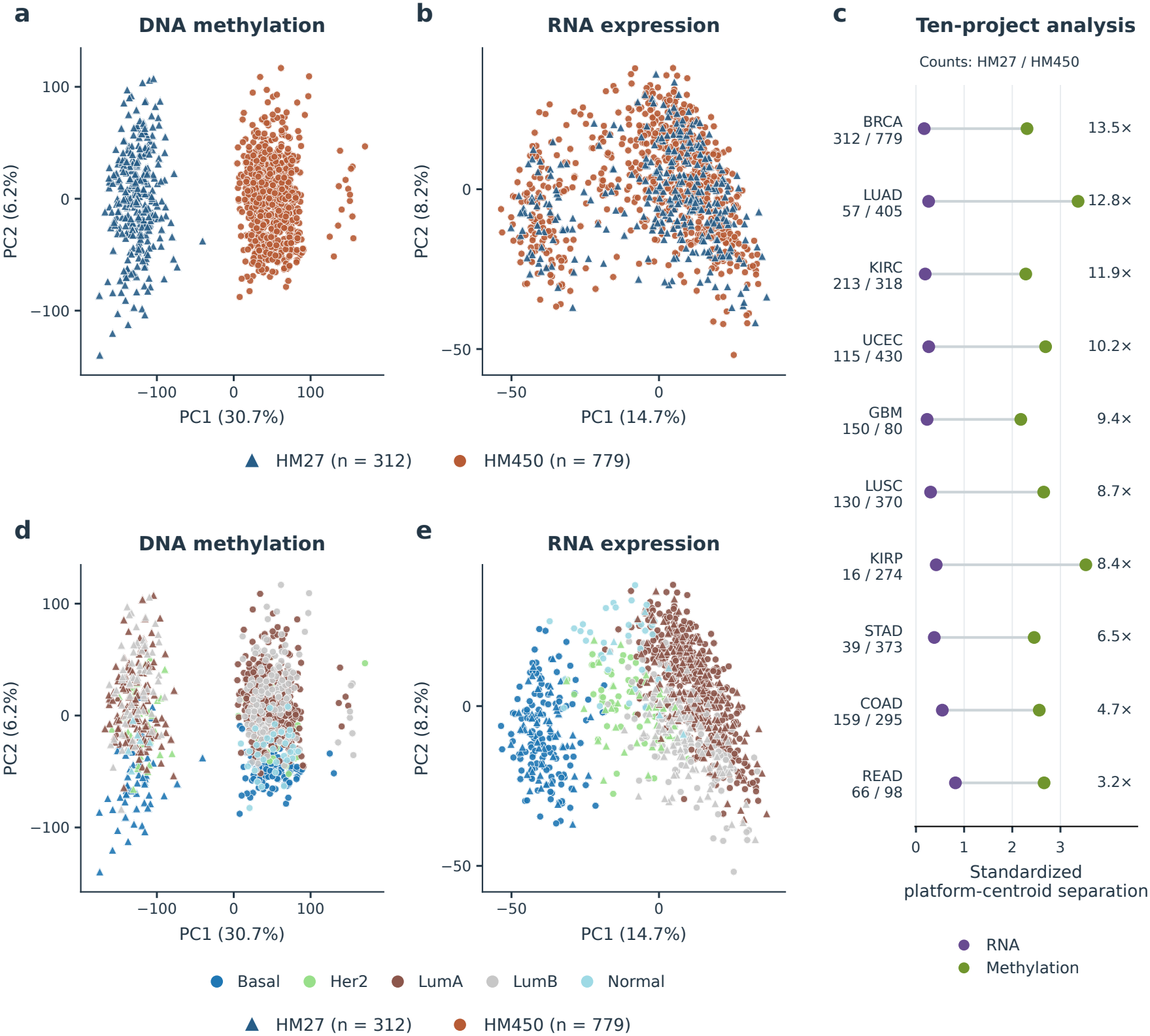
Technical platform asymmetry across paired TCGA multi-omic profiles. The Cancer Genome Atlas (TCGA) paired profiles are shown. **a**,**b**, Uncorrected DNA-methylation (**a**) and RNA-expression (**b**) principal component analysis (PCA) scores for the same 1,091 BRCA tumours, coloured by methylation-array generation: HumanMethylation27 (HM27), triangles (*n* = 312), and HumanMethylation450 (HM450), circles (*n* = 779). Axis percentages denote variance explained in each modality. **c**, Euclidean platform-centroid separation normalized by pooled within-platform dispersion in ten-PC representations of paired profiles across ten TCGA projects. Counts give HM27/HM450 sample numbers. Right-hand annotations give methylation-to-RNA separation ratios (3.2*×*–13.5*×*). **d**,**e**, The identical methylation (**d**) and RNA (**e**) coordinates recoloured by breast-cancer subtype, retaining platform shapes. Axis limits match **a** and **b**, respectively. Patient plots display two PCs. The quantitative analysis uses ten PCs. Subtype labels annotate the plots and do not enter correction fitting.

We therefore used RNA expression as the anchor and DNA methylation as the target for the primary correction benchmark. Seven standard cohorts were included: BRCA, gastrointestinal cancer (GI), UCEC, KIRC, LUSC, LUAD and KIRP (**Supplementary Table 1**). GI combined colon and stomach cancers (COAD and STAD). GBM and READ carried RNA-workflow differences and were excluded from this primary benchmark. GBM was examined separately with the modality roles reversed **(see Methods)**.

Across the seven cohorts, ten methylation principal components (PCs) retained over half of total CpG variation (52.9–60.3%) and captured almost all platform-centroid displacement (*>* 99.9%, **see Methods**). The platform effect dominated the leading component: PC1 alone distinguished the arrays perfectly (Batch AUC 1.00), and platform explained about half of the retained methylation variance (mean Batch EV 51.3%). In paired RNA, the same patient grouping produced much weaker separation (mean Batch AUC 0.73, Batch EV 1.0%). This establishes a critical benchmark for harmonization: when technical groups differ in anchor-supported biological composition, removing that biological separation to force Batch AUC to 0.50 constitutes overcorrection. Representation sensitivity is reported in **Supplementary Table 14**.

### RNA predicts one-third of within-platform methylation variation

Paired RNA predicted about one-third of the within-platform methylation variation represented by ten PCs across the seven standard cohorts (mean out-of-fold *R*^2^ = 0.35). Randomizing RNA–methylation pairings removed this predictability (*R*^2^ = −0.02, **Figure 2**). A multivariate Ridge regression map, termed the cross-modal bridge, captured most of this shared structure with about six predictive directions, providing a compact reference for correction **(see Methods)**.

**Figure 2.**
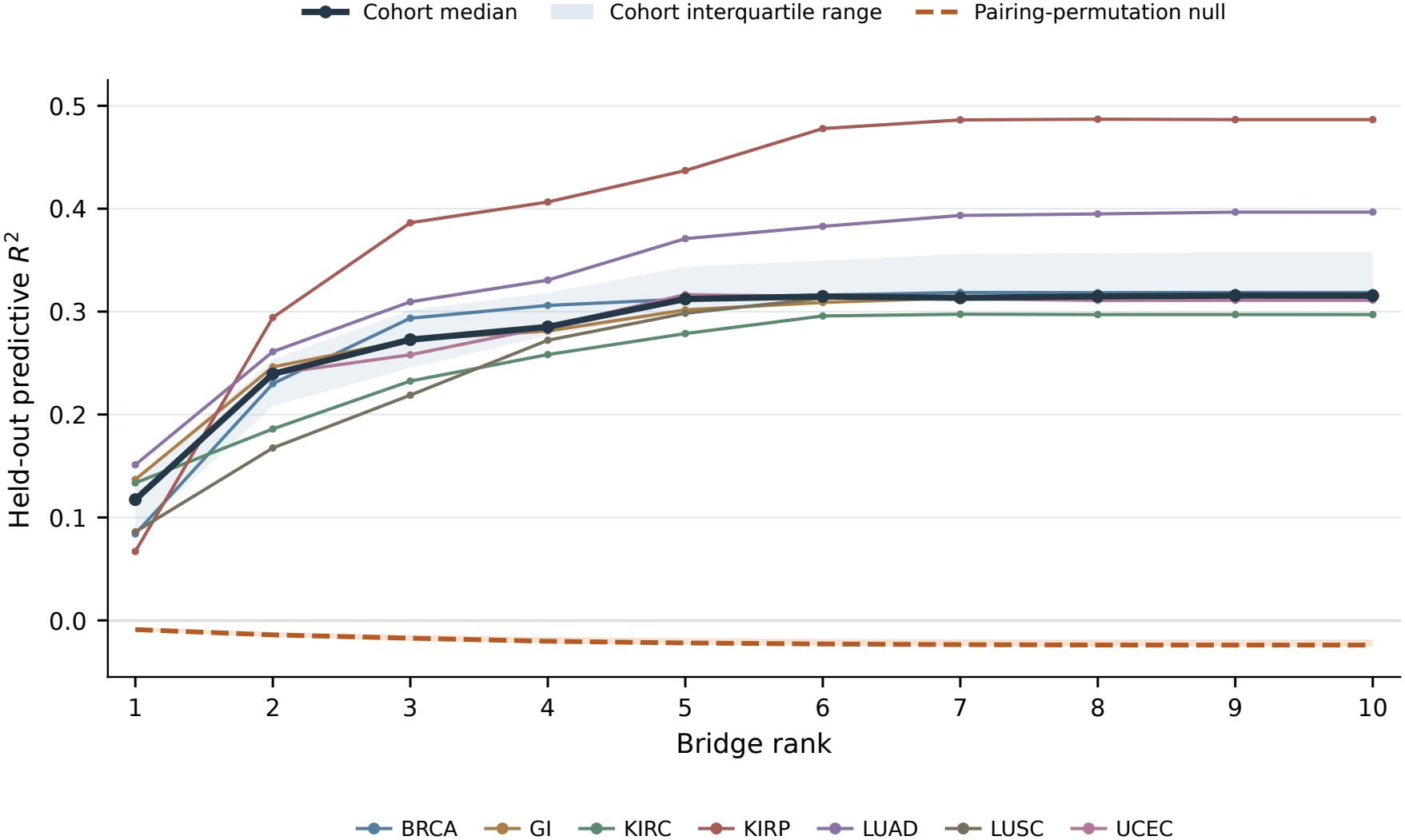
RNA predicts a reproducible low-dimensional methylation subspace. Heldout predictive *R*^2^ for methylation scores from paired RNA across bridge ranks 1–10 in all seven standard cohorts. Thin coloured curves show individual cohorts. The thick dark curve and pale blue band show their median and interquartile range. The dashed orange curve and orange band show the median and interquartile range of cohort-specific means from 200 within-platform pairing permutations. Bands describe between-cohort dispersion. At rank ten, mean paired *R*^2^ was 0.35, compared with −0.02 for the pairing null. All curves use fixed ten-PC representations and matched validation folds **(see Methods)**. Model-capacity comparisons are reported in **Supplementary Figure 1** and **Supplementary Table 5**.

Notably, increasing model complexity offered negligible benefit: polynomial and nonlinear alternatives added little predictive power, and the selected nonlinear models generally operated near their linear regime (**Supplementary Figure 1, Supplementary Table 5**). A simple linear bridge therefore captured the shared structure needed to guide global correction.

Technical distortion can also differ between biological states [4, 5]. Local sample relationships agreed across assays in every cohort, supporting the use of RNA neighbourhoods to guide state-specific correction (mean distance-profile Spearman *ρ* = 0.39, **Supplementary Figure 2**). Five unsupervised RNA neighbourhoods also recovered coherent biology (64% post-hoc subtype purity, **Supplementary Figure 3**). Because categorical labels discretize continuous molecular states, soft memberships allow tumours to contribute across neighbouring states [15, 16]. These global and local correspondences provide the biological reference for the three-stage correction below.

### Cross-modal anchoring enables correction with full feature-level output

Paired RNA provides the reference for distinguishing supported biological differences from technical distortion. The bridge learns the RNA–methylation relationship (**Figure 3a**) and maps RNA into the target methylation space (**Figure 3b**). Three stages then address distinct technical effects: *global translation* removes excess platform-centroid shifts (**Figure 3c**), *global scaling* contracts excess platform spread (**Figure 3d**), and *local translation* uses RNA-defined neighbourhoods (**Figure 3e**) to align state-specific residual offsets (**Figure 3f** ). Reliability and sample-support checks restrict each stage to supported adjustments **(see Methods)**.

**Figure 3.**
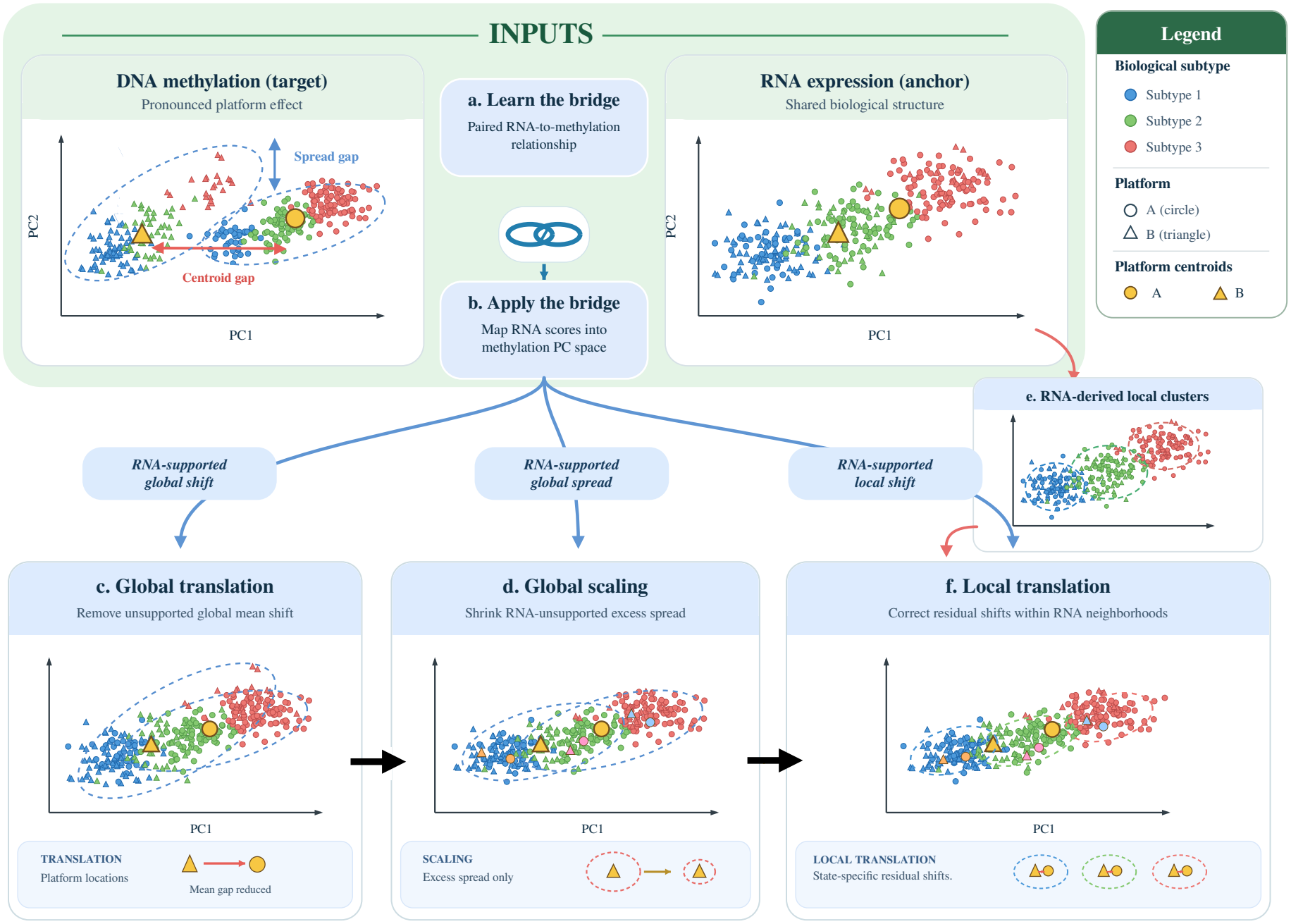
Schematic of the three-stage anchor-guided batch-correction method. **a**,**b**, A bridge learns the paired RNA-to-methylation relationship (**a**) and maps RNA scores into methylation principal component (PC) space (**b**), providing a biological reference for correction. **c**,**d**, Global translation removes the platform-centroid shift not explained by the anchor (**c**). Global scaling contracts excess platform spread (**d**). **e**,**f**, Soft RNA-derived neighbourhoods (**e**) guide local translation of residual state-specific platform offsets (**f** ). Colours denote illustrative biological subtypes, shapes denote platforms, and gold symbols denote platform centroids. The points and contours are schematic.

TCGA subtype annotations were used to evaluate biological preservation. Because the method implicitly learns biological structure from the paired anchor, this protection is expected to extend to continuous and unannotated biology shared across modalities. No biological labels were supplied to the correction.

The correction also retains the full CpG profiles needed for downstream discovery. We add the projected score displacement to each original M-value profile, preserving all variation orthogonal to the corrected subspace to machine precision (**Supplementary Table 7**). This lifting step carries the score corrections back to individual loci without discarding the remaining methylation variation **(see Methods)**. Correction makes full use of the paired cohort, requiring no separate training dataset. The complete pipeline processed the largest cohort, comprising 1,091 breast tumours, in approximately three seconds on a single CPU core using 1.38 GB of peak memory **(see Methods)**.

### Anchor-guided correction reduces platform effects across TCGA cohorts

We benchmarked anchor-guided correction against empirical-Bayes ComBat. *Direct normalization* matches platform means and standard deviations without using the paired anchor. Its performance was comparable to ComBat in standard cohorts, but it failed to preserve subtype structure under confounding (**Supplementary Tables 6a–b, see Methods**).

Across the seven standard cohorts, anchor-guided correction removed nearly all platform-associated variance (mean Batch EV dropping from 51.3% to 0.4%) and brought platform separability close to the paired RNA reference (mean Batch AUC 0.67 versus 0.73 in RNA, **Figure 4** and **Supplementary Figure 4**). ComBat reduced Batch AUC to 0.42, well below the paired RNA reference, suggesting over-alignment. Both methods comparably strengthened molecular subtype structure and made same-subtype tumours easier to identify across arrays than before correction (**Figure 4** and **Supplementary Table 6a**).

**Figure 4.**
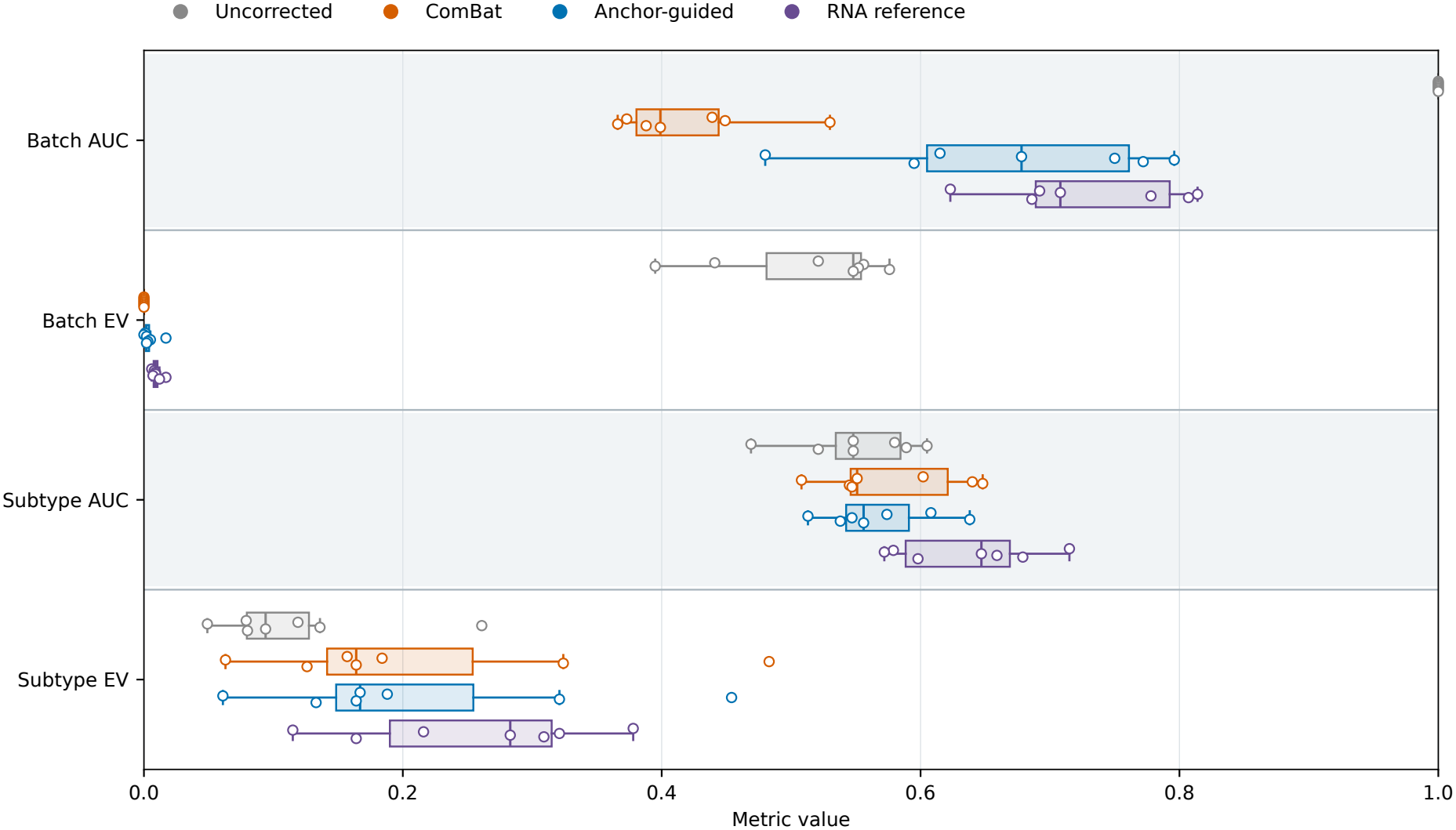
Harmonization performance across seven standard TCGA cohorts. Cohortlevel performance across BRCA, GI, KIRC, KIRP, LUAD, LUSC, and UCEC for uncorrected methylation (grey), empirical-Bayes ComBat (orange), anchor-guided correction (blue), and the paired RNA reference (purple), evaluated by platform predictability (Batch area under the receiver operating characteristic curve, AUC), platform explained variance (Batch EV), crossplatform biological retrieval (Subtype AUC), and subtype explained variance (Subtype EV). Horizontal bands separate the four metrics, each plotted on the same 0–1 scale. Paired RNA-anchor separability (mean Batch AUC 0.73) provides a compositional reference for interpreting platform mixing alongside the biological preservation metrics. Each open circle represents one cohort (*n* = 7). Boxes show interquartile ranges with median lines, and whiskers extend to the most extreme values within 1.5 interquartile ranges of the box. Correction is fitted to the full cohort before classifier cross-validation.

Each stage addressed a different part of the platform effect. Global translation removed the dominant platform shift, global scaling improved local mixing, and local translation resolved state-specific residual offsets (**Supplementary Results**). Neighbourhoods with skewed platform composition became less frequent after scaling: the *k*-nearest-neighbour batch-effect test (kBET) rejection fraction fell from 0.47 to 0.29 **(see Methods)**. The next benchmarks test whether these supported adjustments protect biology when platform composition becomes unequal.

### Cross-modal anchoring resolves batch–biology confounding

We next tested three BRCA benchmarks with progressively less biological overlap between platforms, keeping the measured profiles and PCA axes fixed (**Supplementary Figure 5, Supplementary Table 2, see Methods**).

In the *partial-confounding* benchmark (`BRCA-PC`), both platforms contained Basal and Luminal A tumours in inverted proportions (*n* = 140 per platform). Anchor-guided correction removed platform separation while retaining the subtype organization expected from RNA (**Figure 5a–d**). Cross-platform subtype retrieval improved (Subtype AUC 0.60 to 0.68), whereas ComBat reduced it to 0.58 and decreased subtype-associated variance (**Figure 5e–h, Supplementary Table 6b**). Local translation provided a substantial final improvement in platform alignment, reducing Batch AUC from 0.93 to 0.69 (**Figure 5e**).

**Figure 5.**
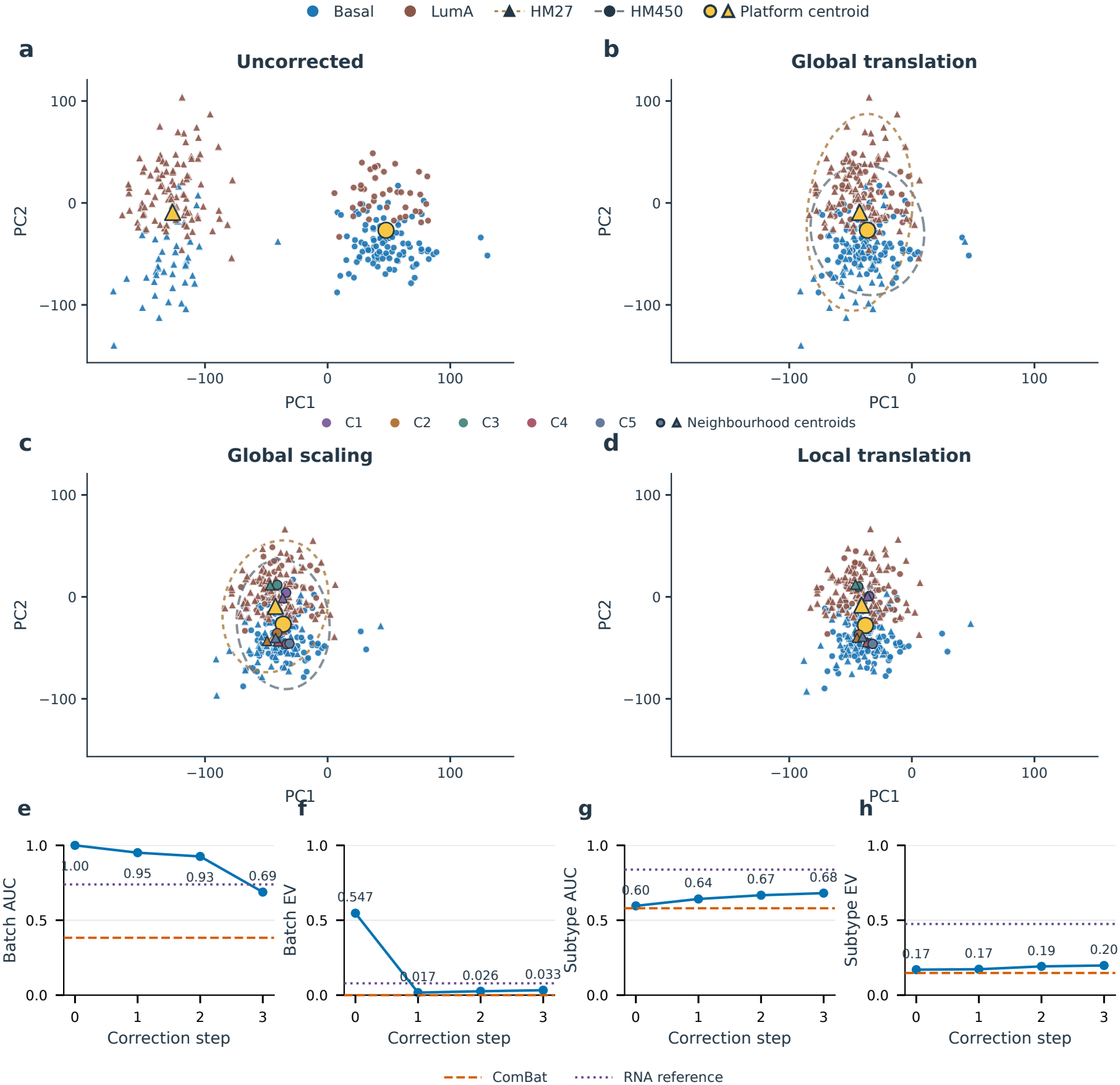
Stage-wise correction in partially confounded breast cancer (BRCA-PC). **a–d**, Observed methylation PC1–PC2 projections in `BRCA-PC` (*n* = 140 per platform with inverted 70%/30% Basal and Luminal A proportions), before correction (**a**), after global translation (**b**), after global scaling (**c**), and after local translation (**d**). Point colours identify subtypes. Circles and triangles denote HM450 and HM27, respectively. Gold symbols show platform centroids. Dashed descriptive 90% Gaussian covariance contours in **b**,**c** show platform spread. In **c**,**d**, smaller coloured circle–triangle pairs joined by segments show platform-specific centroids within the five soft RNA-defined neighbourhoods, calculated using the saved soft memberships. All four panels retain the same PCA axes and coordinate limits. **e–h**, Corresponding ten-PC Batch AUC (**e**), Batch EV (**f** ), cross-platform Subtype AUC (**g**), and Subtype EV (**h**). AUC denotes area under the receiver operating characteristic curve, and EV explained variance. Stages are 0, uncorrected; 1, global translation; 2, global scaling; and 3, local translation. Dashed orange lines show ComBat values. Dotted purple lines show the paired RNA reference for each metric.

The *imbalance support* benchmark (`BRCA-IS`) made this challenge sharper: HM27 contained only Basal tumours (*n* = 55), whereas HM450 contained all five breast cancer subtypes (*n* = 779, **Figure 6a**). Anchor-guided correction aligned the HM27 tumours with the shared HM450 Basal region and improved cross-platform retrieval (Subtype AUC 0.64 to 0.72, **Figure 6b**). ComBat aligned the global platform centroids, placing Basal tumours among mixed-subtype profiles and reducing retrieval to 0.37 (**Figure 6c**). The higher residual Batch AUC after anchoring (0.88 versus 0.44 for ComBat) accompanied better biological preservation (**Supplementary Table 6b**).

**Figure 6.**
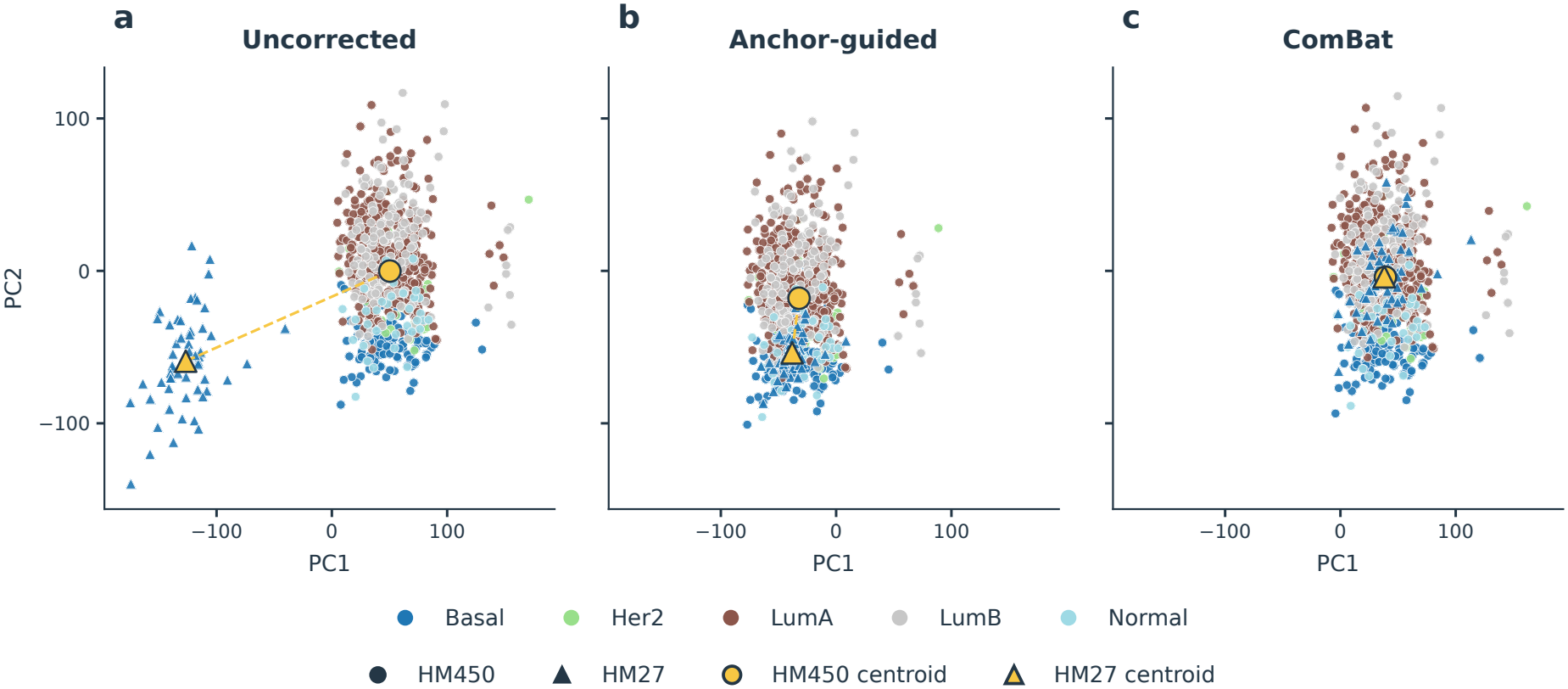
Cross-modal anchoring preserves subtype structure under imbalance support (BRCA-IS). Methylation PC1–PC2 projections before correction (**a**), after three-stage anchor-guided correction (**b**), and after ComBat (**c**) in `BRCA-IS`, where the HM27 array contains exclusively Basal tumours (*n* = 55, triangles) and the HM450 array spans all five PAM50 breast cancer subtypes (*n* = 779, circles). Anchor-guided correction moves the HM27 Basal tumours into the RNA-supported HM450 Basal region while preserving the non-zero compositional offset between global platform centroids (gold symbols). ComBat aligns the global centroids, placing HM27 Basal tumours among mixed-subtype HM450 profiles. All panels retain the same PCA axes and coordinate limits. In the *total-confounding* benchmark (`BRCA-TC`), HM27 contained Luminal A tumours and HM450 contained Basal tumours, with no subtype overlap (*n* = 134 each). Subtype retrieval across platforms is undefined here, so we measured platform/subtype separation **(see Methods)**. ComBat collapsed the biological distinction (separation AUC 1.00 to 0.40). Anchor-guided correction preserved it (AUC 1.00) by rejecting local alignment where biological overlap was absent (**Supplementary Table 6b**). These benchmarks show that supported alignment can recover shared biology without forcing unmatched biological states together.

In the *total-confounding* benchmark (`BRCA-TC`), HM27 contained Luminal A tumours and HM450 contained Basal tumours, with no subtype overlap (*n* = 134 each). Subtype retrieval across platforms is undefined here, so we measured platform/subtype separation **(see Methods)**. ComBat collapsed the biological distinction (separation AUC 1.00 to 0.40). Anchor-guided correction preserved it (AUC 1.00) by rejecting local alignment where biological overlap was absent (**Supplementary Table 6b**). These benchmarks show that supported alignment can recover shared biology without forcing unmatched biological states together.

### Null experiments confirm biological preservation and controlled false-positive rates

Correction should remain restrained when batch labels do not represent a technical difference. Across 50 random partitions in each of three single-platform cohorts, anchor-guided correction preserved subtype retrieval on average and kept artificial-group separation close to chance (mean Batch AUC 0.47, compared with 0.44 for ComBat and 0.35 for Direct normalization, **Supplementary Figure 6, Supplementary Table 8**). The representative PCA example shows the limited change in sample relationships, including a spread adjustment without centroid movement (**Supplementary Figure 6**).

False-positive rates remained at expected levels after lifting. Across 180 genome-wide null scans, the fraction of nominally significant CpGs matched its expected level (4.9% at *p <* 0.05), and 97.8% of scans produced no false discoveries after multiple-testing correction (**Supplementary Table 9, see Methods**). These results support testing biological discovery on the lifted CpG profiles.

### Cross-modal anchoring improves CpG discovery under imbalance support

We tested whether preservation of sample relationships translated into preservation of individual methylation effects. Across four established tumour contrasts [17–19], lifted profiles retained strong cross-platform agreement in the three high-signal comparisons (concordance correlation coefficient, CCC, 0.84–0.93). In the subtler UCEC MSI–POLE contrast, concordance exceeded ComBat (CCC 0.25 versus 0.22, **Figure 7a, Supplementary Table 10**).

**Figure 7.**
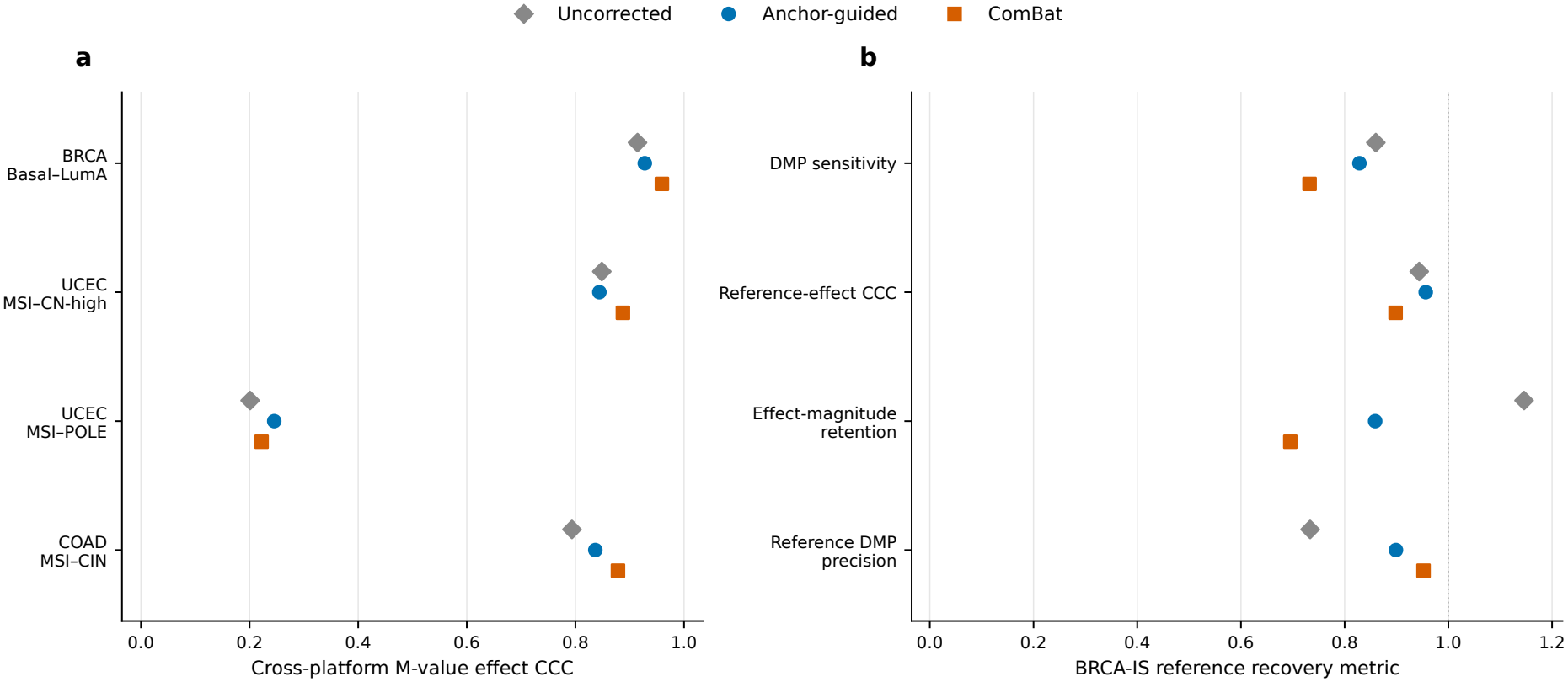
Cross-platform locus-level concordance and CpG discovery under imbalance. **a**, Lin’s concordance correlation coefficient (CCC) for CpG-level differential methylation M-value effects across four prespecified tumour contrasts (BRCA Basal–Luminal A, UCEC MSI–CN-high, UCEC MSI–POLE, and COAD MSI–CIN) before correction (grey diamonds), after anchorguided correction (blue circles), and after ComBat (orange squares). MSI denotes microsatellite instability, CN-high high copy-number alteration, POLE polymerase-epsilon-mutant tumours, and CIN chromosomal instability. **b**, Patient-disjoint recovery of 5,346 within-HM450 Basal-versus-Luminal-A reference differentially methylated probes (DMPs, false discovery rate, FDR, below 0.05 and |Δ*β*| ≥ 0.05) in the imbalance support case (`BRCA-IS`), comparing same-direction DMP sensitivity, beta-scale reference-effect CCC, median effect-magnitude retention, and reference DMP precision between lifted anchor-guided M-values and ComBat. Grey diamonds show the uncorrected baseline. Retention is the median absolute corrected-to-reference effect ratio: 1 denotes matched magnitudes, values below 1 attenuation, and values above 1 amplification. Correction was fitted to the full benchmark cohort before patient-disjoint downstream effect estimation.

Recovery of differentially methylated probes (DMPs) was comparable under partial confounding in `BRCA-PC` (approximately 80% for both methods). Against an independent 1,741-CpG signature from GSE75067 [20], both methods recovered over 98% of markers in `BRCA-IS`, with anchorguided correction retaining effect magnitudes closer to the external reference (**Supplementary Table 11, see Methods**).

Under imbalance support, anchor-guided correction improved DMP recovery compared with ComBat. We tested `BRCA-IS` against 5,346 reference DMPs defined in independent patients from the same HM450 cohort **(see Methods)**. Anchor-guided correction recovered more reference DMPs than ComBat (82.8% versus 73.2% sensitivity), with stronger effect-size concordance (CCC 0.96 versus 0.90, **Figure 7b**). It also retained more of their effect magnitudes (85.9% versus 69.5%). Uncorrected data had higher sensitivity (86.0%) but lower precision (73.3%) than anchor-guided correction (89.9% precision), which achieved the highest effect-size concordance (**Supplementary Table 11**). The gain therefore reflects a better balance between DMP discovery and preservation of methylation effects.

### Methylation can guide RNA workflow correction

We reversed the assay roles in glioblastoma (GBM), using methylation to anchor RNA workflow correction. Patients were analysed separately within the HM27 and HM450 subsets, where the workflows had broadly overlapping subtype composition. Simple corrections based on externally selected housekeeping (HK) genes [21] failed to reduce workflow separation. We therefore tested a multivariate HK approach allowing different responses across RNA principal components (`HK-PC`, **see Methods**). Anchor-guided correction reduced workflow-associated variance by approximately 97% in both subsets. Residual RNA separation remained above the methylation anchor reference, while HK-PC and ComBat reduced separation below it. All three methods increased subtype-associated variance, with the largest gains from anchor-guided correction and ComBat (**Supplementary Table 13**).

### Anchoring tolerates independent noise and preserves private biology

Finally, simulations separated biological and technical components under known ground truth. Across 30 paired seeds, we varied anchor contamination and the amount of biology shared between assays (**Supplementary Table 4, see Methods**).

Independent technical noise in the RNA anchor left methylation correction and biological preservation virtually unchanged across the tested range (**Supplementary Results**). Shared technical contamination behaved differently: as the same artefact entered both assays, it increasingly remained after correction (**Supplementary Figure 7**). As expected, these shared technical effects led to under-correction, while shared biology remained protected throughout these tests. Biology unique to methylation was also retained. With just 5% of target biological variance shared with RNA, correction removed approximately 71% of the technical effect and preserved 92% of the shared component (**Supplementary Figure 8a–d**). Preservation of the methylationprivate component remained approximately 95% across the tested range (**Supplementary Figure 8c**,**d**). Thus, even limited biological overlap can support effective correction while preserving both shared and methylation-private biology.

## Discussion

Standard unimodal batch-correction methods such as ComBat are designed to remove all systematic differences between batches [6–8]. While this works well when batches have the same biological composition, it creates a major blind spot whenever sample composition differs across technical groups [4, 5, 10]. Because a single assay cannot tell a genuine biological difference apart from a technical offset, unimodal methods systematically erase biological signal when batches are imbalanced [6, 7, 10]. Worse, standard batch-mixing metrics cannot warn the user when overcorrection happens [6, 7]. Making batches indistinguishable looks the same whether a method removed technical noise or wiped out a real biological difference between batches [6, 7]. The breast cancer benchmarks expose this exact problem across increasing levels of compositional imbalance (**Figures 5–6, Supplementary Table 6b**). Rather than forcing batches together blindly, anchor-guided correction uses the paired modality to adapt to the setting. It aligns batches fully when their biology overlaps, aligns only shared subpopulations when composition is skewed, stops local alignment when batches share no biology, and leaves the data untouched when no real batch effect is present (**Supplementary Figure 6, Supplementary Table 8**).

A major practical advantage of this approach is that it requires no clinical annotations or predefined covariates. Although unimodal methods such as ComBat allow users to include covariates to protect known groups, any biological variation that is not explicitly supplied as a covariate is treated as a technical batch effect and removed [8, 10]. In practice, this leaves unannotated subtypes, plastic cell states, and unmeasured continuous gradients such as tumour purity vulnerable to erasure [8, 10]. By contrast, a paired molecular anchor acts as an implicit, continuous reference that does this work automatically for the user. Through the linear bridge and local anchor neighbourhoods, the anchor captures shared biological variation in the cohort and protects that structure without requiring prior labels **(see Methods)**.

Correcting batch effects in the target assay’s native coordinate system also separates our method from existing multi-omic tools. Factor models such as MOFA+ [11,12] combine modalities into an abstract latent space for exploratory analysis, while cross-omic pipelines such as MultiBaC [13] impute missing layers before applying standard correction. Neither approach resolves batch– biology confounding while returning the affected assay in its original feature space. Conceptually, our method separates variation in the target assay into three parts. The first is shared biology predicted by the anchor, which is directly protected. The remaining variation is specific to the target assay and splits into modality-private biology, which is protected by our statistical gates and orthogonal feature lifting, and technical batch effects, which are isolated and removed. Because the lifting step maps low-rank adjustments back to the original features without losing orthogonal variation, users receive corrected feature-level profiles, such as CpG M-values or gene expression, ready for any standard downstream task with controlled false-positive rates (**Supplementary Tables 7 and 9**). In simulations, this design retained roughly 95% of modalityprivate biology (**Supplementary Figure 8c**,**d**). In the imbalanced breast cancer benchmark, it directly improved differential methylation discovery, whereas overcorrection by ComBat damaged true signal recovery more than leaving the batch effect uncorrected (**Figure 7b, Supplementary Table 11**).

Finally, cross-modal anchoring is a portable concept that is not tied to a specific pair of technologies. Reversing the modality roles in glioblastoma, where DNA methylation anchored RNA-sequencing workflow correction, shows that the method works in either direction and outperforms simple housekeeping-gene adjustments [9,21] (**Supplementary Table 13**). Applying the method relies on three testable assumptions: the assays must be paired across samples, they must share a reproducible biological subspace, and the anchor must be relatively stable across the target’s batches. As our bridge analyses and simulations show, a simple linear mapping captures this shared space even when biological overlap is modest, with stable correction across the tested range from 5% shared biological variance upwards (**Figure 2, Supplementary Figure 8**). Moreover, the anchor does not need to be completely free of batch effects; independent technical noise in the anchor is filtered out by the bridge, while technical artefacts shared across both assays lead safely to conservative under-correction rather than biological erasure (**Supplementary Figure 7**). Looking ahead, this framework can be naturally extended in several ways. Future work could combine multiple stable layers into joint anchors, support semi-anchored designs where only a subset of samples is paired across modalities, or integrate the cross-modal anchor directly into established unimodal estimators and control-gene methods. Testing these extensions across other molecular layers, such as proteomics and chromatin accessibility, and scaling them to single-cell multi-omic atlases will broaden the reach of cross-modal anchoring for confounded biological studies.

## Methods

### TCGA data and study design

Paired gene-expression and DNA-methylation profiles were obtained from the National Cancer Institute Genomic Data Commons (GDC) [1, 22]. GDC Spliced Transcripts Alignment to a Reference (STAR) gene counts [23] and processed SeSAMe level-3 methylation beta values [24] were joined deterministically by the 12-character participant-level TCGA barcode and restricted to Primary Tumor specimens measured in both modalities (Supplementary Methods and Supplementary Table 1). Throughout the primary pan-cancer analyses, the technical batch variable was array generation, comparing the Illumina Infinium HumanMethylation27 (HM27) and HumanMethylation450 (HM450) BeadChips across their shared probe intersection. Before anchor selection, PCA diagnostics were inspected against sequencing workflow, processing plate and centre metadata to distinguish assay-wide separation from localized technical groups (Supplementary Methods).

The cross-modal technical asymmetry analysis encompassed ten TCGA projects with paired samples spanning both methylation arrays: BRCA, COAD, GBM, KIRC, KIRP, LUAD, LUSC, READ, STAD, and UCEC (Supplementary Table 1). For primary harmonization benchmarks where RNA-seq served as the anchor and DNA methylation as the target, COAD and STAD were pooled after project-specific preprocessing into a gastrointestinal (GI) cohort, yielding seven standard evaluation cohorts: BRCA (*n* = 1, 091), GI (*n* = 866), KIRC (*n* = 531), KIRP (*n* = 290), LUAD (*n* = 514, with 462 on HM27/HM450), LUSC (*n* = 500), and UCEC (*n* = 545). Glioblastoma (GBM, *n* = 230) was reserved for a role-reversed workflow-control analysis because its RNA-seq profiles carried a distinct sequencing-workflow batch effect, allowing DNA methylation to be evaluated as the anchor modality and RNA-seq as the target. READ (*n* = 164) was excluded from the primary RNA-anchored benchmark because its RNA-sequencing workflow co-varied with methylation-array generation and remained associated with RNA structure after subtype adjustment (Supplementary Methods).

### Modality preprocessing and principal component models

RNA-seq genes were retained when expression exceeded 1 count per million (CPM) in at least max(5, ⌈0.1*n*⌉) samples, normalized by the trimmed mean of M-values (TMM), transformed to log_2_(CPM + 1), restricted to the 3,000 most variable genes, and standardized to zero mean and unit variance prior to principal component analysis (PCA) [25, 26]. In DNA methylation, common-array CpG probes with more than 5% missingness were excluded, remaining missing values were imputed using uniform 15-nearest-neighbour imputation across samples [27], and sex-chromosome probes were removed [28]. Bounded methylation beta values (*β*) were converted to logit M-values via *M* = log_2_[(*β* +10^−6^)*/*(1−*β* +10^−6^)] [29], followed by removal of zero-variance probes and feature-wise standardization (21, 635–22, 814 retained CpGs across primary cohorts).

Separate PCA models were fitted to each modality to concentrate coherent multivariate variation, regularize the cross-modal bridge, and provide orthogonal coordinates for interpretable correction [26]. The primary representation retained *r*_RNA_ = *r*_METH_ = 10 principal components per modality (*Z*^RNA^, *Z*^METH^ ∈ ℝ ^*n*×10^). For the centroid shift Δ across all retained CpGs in the standardized M-value matrix and leading loading rows *V*_10_ ∈ ℝ ^10×*D*^, the captured squared platform-centroid separation was

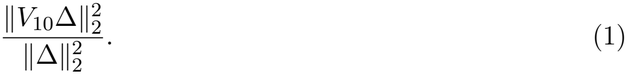

Representation sensitivity was assessed in all seven standard cohorts and three BRCA confounding benchmarks at ten and 50 components per modality, using the same patients, comparator methods and three-stage correction settings (Supplementary Table 14). The RNA neighbourhood representation used the selected rank, with the number of neighbourhoods and all reliability, support and permutation thresholds held fixed. Metrics were evaluated within each retained-score representation to assess score-space sensitivity. Additional paired-platform mapping experiments in AML and LUAD compared ten- and 50-PC linear correction spaces (Supplementary Methods).

### Cross-modal bridge and nonlinear capacity benchmark

Let *Z*^RNA^ and *Z*^METH^ ∈ ℝ ^*n*×10^ denote the paired ten-component score matrices pooled across platforms *b*_*i*_ ∈ {0, 1}. The cross-modal bridge used for correction was fitted once per cohort to the uncentred pooled scores via multivariate Ridge regression [30]:

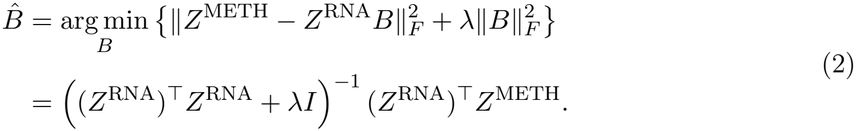

The trace-normalized penalty was

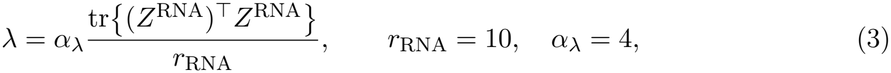

with settings listed in Supplementary Table 3. Trace normalization scales the penalty to the mean anchor-PC sum of squares, allowing the dimensionless multiplier *α*_*λ*_ to be shared across cohorts. The linear system was solved directly without explicit matrix inversion to yield the anchor-predicted target scores 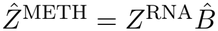.

To quantify reproducible shared biology independently of platform shifts, out-of-fold bridge predictability was evaluated in all seven standard cohorts across five platform-stratified folds (with platform means estimated strictly within training folds) over bridge ranks 1–10, benchmarked against 200 within-platform training-sample pairing permutations per cohort. Rank curves used a fixed Ridge penalty *λ* = 1 and the fixed ten-PC representations. LUAD was restricted to its 462 HM27/HM450 patients. Permuted bridges were evaluated against unchanged held-out target profiles using the same folds as the paired bridge (Supplementary Methods). To test whether nonlinear mappings captured additional shared structure, linear Ridge was compared against degree-two polynomial Ridge (including linear, squared, and pairwise interaction terms) and radial basis function (RBF) Kernel Ridge [31] across identical five-fold outer and three-fold inner platform-stratified cross-validation splits using pooled out-of-fold Frobenius *R*^2^ and 2,000 case-bootstrap 95% confidence intervals (Supplementary Methods and Supplementary Table 5).

### Three-stage anchor-guided correction

Three sequential corrections were applied in the fixed target PCA space, reusing the same pooled bridge 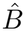 and identical hyperparameter thresholds across all cohorts (Supplementary Table 3). Let 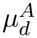 denote the centroid of platform *d* ∈ {0, 1} in modality *A*, and 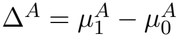 denote the inter-platform centroid gap.

**Stage 1: Global translation**. Global translation compares the observed target centroid gap Δ^METH^ against the anchor-predicted gap 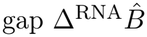. For discrepancy vector 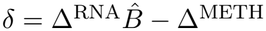 and binary PC-wise gate vector *g* ∈ {0, 1}^10^, the translated scores are:

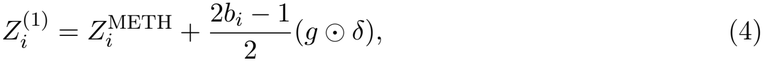

where ⊙ denotes element-wise multiplication. Admitted components receive symmetric, opposite shifts that preserve the midpoint of the two platform centroids. Gate *g*_*j*_ = 1 is admitted only when three criteria hold simultaneously: (i) component *j* achieves non-negative five-fold platform-centred cross-validated *R*^2^; (ii) the 95% interval of *δ*_*j*_ across 30 within-platform paired bootstrap resamples excludes zero; and (iii) the proposed shift reduces the gap magnitude 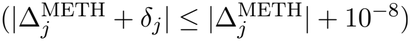 without overshooting or reversing the sign of 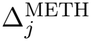.

**Stage 2: Global scaling**. Global scaling compares robust within-platform variances after global translation 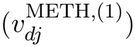 against the corresponding variances of the bridge prediction 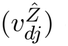 computed on the same outlier-fenced core (excluding observations beyond 1.5 times the interquartile range from the median, capped at 5% per platform and PC). For component *j*, the residual log-spread ratio and contraction factor are:

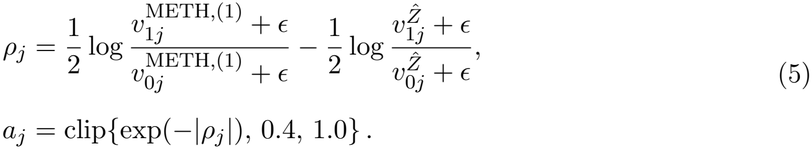

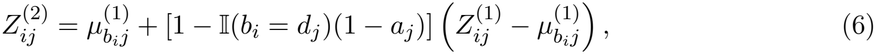

where *d*_*j*_ = 1 if *ρ*_*j*_ *>* 0 and *d*_*j*_ = 0 otherwise. Thus, only the platform exhibiting wider dispersion than predicted by the anchor is contracted about its own centroid, preventing artificial variance inflation. The stage requires at least 15 samples per platform and a root-mean-square residual spread across PCs exceeding the 95th percentile of 50 platform-label permutations. Otherwise *a*_*j*_ = 1.

**Stage 3: Local translation**. We fit *K* = 5 unsupervised *K*-means centres [32] to pooled anchor scores, with 30 initializations. The cluster-count sensitivity analysis is shown in Supplementary Figure 3. Squared distances 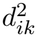 to centre *k* are converted to soft memberships using

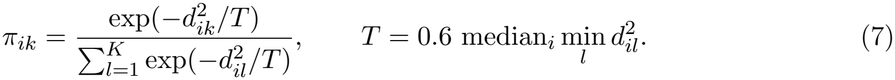

Effective platform support, membership-weighted platform centroids, and local gaps are defined by

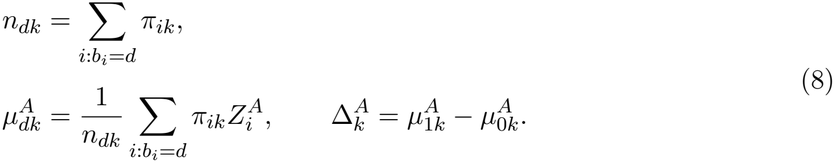

The local residual and support weights are

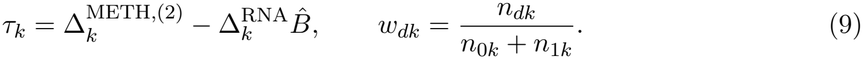

For gated residual vectors *h*_*k*_, the final score state is

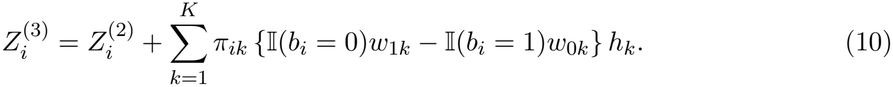

Neighbourhood *k* is admitted only when both platforms contribute effective support *n*_*dk*_ ≥ 10 and ∥*τ*_*k*_∥_2_ exceeds the 95th percentile of 50 platform-label permutations. Admitted vectors *h*_*k*_ are capped at three times the smaller within-platform membership-weighted standard deviation projected along *τ*_*k*_*/*∥*τ*_*k*_∥_2_. An optional fourth exploratory stage for contracting isolated local tail outliers with a *k*-nearest-neighbour local-density filter is formulated in the Supplementary Methods but excluded from the primary three-stage endpoint.

### Cycle-consistent M-value lifting and computational execution

To return corrected PCA scores 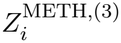 to CpG-level M-values without truncating highdimensional variation outside the ten-PC subspace, only the low-rank score displacement 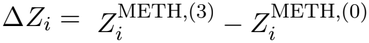 is lifted through the fixed orthonormal PCA loading rows *V* ∈ ℝ^10×*D*^:

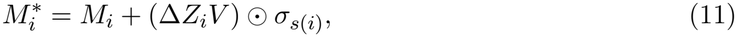

where *M*_*i*_ ∈ ℝ^*D*^ is the original M-value vector and *σ*_*s*(*i*)_ ∈ ℝ^*D*^ contains the feature standard deviations of the preprocessing model for sample *i*’s source project. Every exported matrix was verified against eight deterministic numerical invariants (tolerances ≤ 2 × 10^−5^ absolute and 10^−6^ relative) confirming exact preservation of the orthogonal feature-space residual 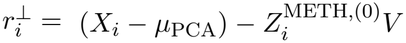 and exact recovery of 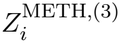 upon reprojection both in float64 and after float32 serialization (Supplementary Methods and Supplementary Table 7).

Computationally, for *n* samples, *D* probes, *r* = 10 PCs, and *K* = 5 neighbourhoods, the bridge and local offsets contain *o* (*r*^2^ + *Kr*) coefficients, with *o* (*nK*) sample memberships. Projection through fixed PCA loadings and lifting each cost *o* (*nDr*), while loading the full feature matrix requires *o* (*nD*) memory. The pooled Ridge bridge is fitted once for correction and reused across stages, with additional fits for cross-validation and resampling gates. All paired patients contribute to correction without requiring a separate training dataset. On a single CPU core for the largest cohort (BRCA, *n* = 1, 091 samples and *D* = 22, 199 probes), the complete pipeline from input M-values and paired RNA profiles through PCA projection, three-stage correction and CpG lifting executed in 2.93–3.06 s with 1.38 GB peak resident memory (Supplementary Methods).

### Evaluation cohorts, comparators, and metrics

Alongside the seven standard TCGA cohorts, three controlled benchmarks were constructed from TCGA-BRCA by subsetting samples while holding molecular profiles and full-cohort PCA axes fixed (Supplementary Table 2): partial confounding (`BRCA-PC`, *n* = 140 per array with inverted 70%/30% Basal and Luminal A proportions), imbalance support (`BRCA-IS`, *n* = 779 HM450 across all five PAM50 subtypes and *n* = 55 HM27 Basal only), and total confounding (`BRCA-TC`, *n* = 134 HM450 Basal and *n* = 134 HM27 Luminal A). All scenarios compared uncorrected methylation, Direct normalization, parametric empirical-Bayes ComBat [8] applied to CpG-level M-values (implemented via `inmoose`/`pycombat_norm` and projected onto the fixed PCA axes), and three-stage anchor-guided correction. For Direct normalization, each platform’s scores were centred and scaled separately along each PC, then mapped to the pooled mean and pooled within-platform standard deviation. This preserves the relative scale of the PCs, while forcing equal platform centroids and spreads without cross-modal information.

Four primary metrics were evaluated on identical score coordinates (Supplementary Table 6 and Supplementary Methods): (i) *Batch AUC*, the mean five-fold stratified logistic-regression ROC AUC for predicting methylation platform; (ii) *Batch EV*, the fraction of total squared score-space variance explained by platform centroids (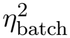); (iii) *Subtype AUC*, the equal-weight mean of subtype-specific cross-platform retrieval ROC AUCs calculated from negative Euclidean distances over subtypes represented on both arrays; and (iv) *Subtype EV*, the fraction of score-space variance explained by molecular subtype centroids (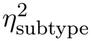 ). Local batch mixing was monitored using a binary implementation of the *k*-nearest-neighbour batch-effect test (kBET) [33]: Pearson’s chi-square test compared the two-platform composition of each 30-neighbour neighbourhood with the global platform proportions, and the rejection fraction used a significance threshold of 0.05.

### Random-label stability and CpG false-positive testing

Stability under artificial null partitions was evaluated across 50 independent random binary partitions (seeds 0–49) in three single-platform HM450 cohorts: PRAD (*n* = 501, split 250*/*251), THCA (*n* = 505, split 252*/*253), and LIHC (*n* = 371, split 186*/*185). The sham batch labels were assigned identically to paired RNA and methylation profiles without altering molecular values, and subtype AUC and artificial-group AUC were recorded after three-stage correction.

False-positive rates in genome-wide tests of lifted M-values were evaluated independently. PRAD, THCA, and LIHC were each partitioned into three synthetic-platform splits, and the complete three-stage correction and M-value lift were fitted independently to each split. Twenty new balanced binary sham phenotypes were then generated per lifted matrix (yielding 180 genome-wide scans across 363, 416–398, 782 CpGs per method arm) and tested via empiricalBayes moderated linear models (`limma`) [34]. False-positive behaviour was assessed using mean _,obs_ *p*-value, nominal *p <* 0.05 fraction, genomic inflation factor 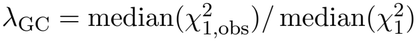, and Benjamini–Hochberg false discoveries (*q <* 0.05) [35].

### Feature-level validation and CpG discovery benchmarks

Downstream CpG-level evaluation comprised four designs using the empirical-Bayes moderated linear models in `limma` [34], implemented in `inmoose`, with Benjamini–Hochberg false discovery rate (FDR) control [35].

#### 1. Cross-platform effect concordance in standard cohorts

Four prespecified contrasts were evaluated over shared probes: BRCA Basal versus Luminal A, UCEC MSI versus copynumber-high, UCEC MSI versus POLE, and COAD MSI versus CIN [17–19]. Moderated M-value effects and beta-scale differences (Δ*β*) were estimated independently within HM27 and HM450. Cross-array agreement was quantified using Lin’s CCC [36] and Spearman correlation of M-value effects, and directional agreement among probes with |Δ*β*| ≥ 0.05 on at least one array. Targeted controls included the *MLH1* promoter and canonical CIMP markers (*RUNX3, CACNA1G, IGF2, NEUROG1*, and *SOCS1* ) [37] (Supplementary Table 10).

#### 2. Patient-disjoint DMP discovery under imbalance support (BRCA-IS)

An internal Basal-versus-Luminal-A reference was defined within HM450, using FDR below 0.05 and |Δ*β*| ≥ 0.05. A fixed reference-definition subset identified 5,346 DMPs. Those patients were excluded from downstream DMP estimation in the evaluation subset. The corrected matrices were fitted to the original full benchmark cohorts before this downstream patient split. Recovery was quantified by same-direction sensitivity, precision, CCC of beta-scale effects against the reference, and the median corrected-to-reference absolute effect ratio (Supplementary Table 11). A secondary analysis used the complete within-HM450 reference before patient splitting.

#### 3. Patient-disjoint DMP discovery under partial confounding (BRCA-PC)

The same fixed 5,346-DMP reference and downstream patient-disjoint protocol were applied to `BRCA-PC`, where both subtypes were represented on both arrays in inverted proportions (Supplementary Table 11).

#### 4. Independent external replication (GSE75067)

An independent HM450 breast-cancer cohort [20] supplied 154 primary tumours with expression-derived PAM50 labels, including 43 Basal and 41 Luminal A tumours. All five subtypes contributed to variance estimation in an M-value model adjusted for age and processing plate. The Basal-minus-Luminal-A contrast defined 1,741 shared CpGs with FDR below 0.05 and absolute adjusted Δ*β* ≥ 0.10, without using TCGA outcomes. This fixed external reference was used to evaluate sensitivity, beta-scale effect CCC and median effect-magnitude retention in corrected `BRCA-IS`, using the target DMP threshold |Δ*β*| ≥ 0.05 and FDR below 0.05 (Supplementary Table 11).

### Role-reversed glioblastoma and housekeeping-gene benchmarks

To test DNA methylation as the anchor for an RNA sequencing-workflow effect, we compared correction methods within the HM27-associated (*n* = 150) and HM450-associated (*n* = 79) TCGA-GBM subsets. Each subset contained the two major RNA workflows. One additional HM450 patient had a third workflow represented by a single sample and was excluded. All methods were fitted to the full eligible subsets.

#### 1. External housekeeping control panel

Housekeeping candidates from the HRT Atlas [21] and literature were screened using external disease annotations and glioblastoma-versus-nontumour expression equivalence in GSE4290 [38]. This defined 1,138 genes, of which 1,112 mapped uniquely to measured RNA genes. TCGA subtype labels and within-cohort variability were not used to select the panel (Supplementary Methods).

#### 2. Housekeeping-based correction

Workflow-specific location (median shift), robust affine (Huber regression) [39], and LOWESS [40] transformations were estimated from the HK genes and applied to the 3,000 RNA features used in the original GBM analysis. For `HK-PC`, the standardized HK expression difference between workflows defined a sample-level technical factor. A separate coefficient was fitted for each of ten RNA PCs, and the fitted factor contribution was subtracted when the HK workflow difference passed its permutation test (Supplementary Methods).

#### 3. Anchor-guided correction, ComBat and evaluation

Ten methylation PCs anchored correction of ten RNA PCs using the fixed global translation, global scaling and local translation settings in Supplementary Table 3. ComBat was applied to the same standardized RNA features and projected into the same RNA PCA coordinates. Batch AUC, Batch EV, Subtype AUC and Subtype EV used the primary definitions above, with RNA workflow replacing methylation platform as the batch label. Subtype AUC measured retrieval across workflows. The paired methylation scores supplied the anchor reference (Supplementary Table 13).

### Behavioural simulations

The multi-omic simulator generated *n* = 400 paired samples across 1, 200 RNA features and 3, 000 methylation CpGs with four balanced biological subtypes, two technical batches, and ten PCs per modality across 30 paired random seeds (seeds 1001–1030, Supplementary Methods and Supplementary Table 4). For modality *d* ∈ {*R, M* }, clean biological matrices were generated as

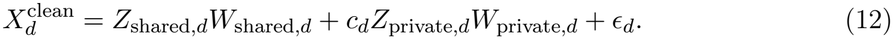

For the shared-biology sweep, with *ϕ* ∈ [0, 1], the target was parameterized as

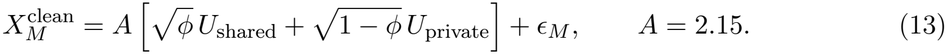

Target methylation technical structure was added as *E*_*M*_ = *s*_*M*_ (2.20*b* + 0.35*u*_*M*_ )*w*_*M*_ for batch indicator *b* ∈ {−0.5, +0.5} and sample-level technical heterogeneity *u*_*M*_ ∼ *N* (0, 1). Holding target methylation and true biological components fixed within each seed, three stress sweeps varied: (i) independent RNA technical contamination (*s*_*R*_ ∈ {0.00, 0.08, 0.13, 0.19, 0.29} realized RMS with *u*_*R*_ ⊥ *u*_*M*_ ); (ii) shared cross-modal technical contamination across a 5 × 5 factorial grid crossing *s*_*R*_ ∈ {0.00, 0.30, 0.75, 1.50, 3.00} with shared technical fraction *f* ∈ {0, 0.25, 0.50, 0.75, 1.00}; and (iii) shared biological fraction *ϕ* ∈ [0%, 100%]. For correction displacement 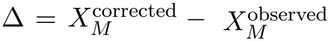 and true technical matrix 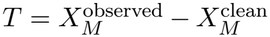, technical removal and preservation of biological component *B* (shared or private) were measured as

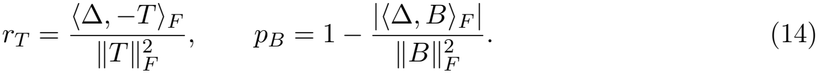

The preservation score measures correction displacement projected onto the known biological component. It is distinct from a fraction of total biological variance retained.

### Statistics and reproducibility

Patients were the biological units of analysis. TCGA analyses used all eligible paired primary tumours under the stated inclusion rules, with cohort sizes and platform composition reported in Supplementary Tables 1 and 2. The principal correction results describe harmonization of the observed cohorts. Correction was fitted before classifier cross-validation, so platform-classifier AUC quantifies residual separability in the corrected cohort. It does not measure correction performance on unseen patients. Bridge capacity was assessed using held-out folds in the fixed preprocessed representation. The GBM RNA-workflow benchmark fitted all correction methods to the full eligible cohorts and evaluated residual separation and subtype structure in those cohorts. Across 50 platform-stratified 80% cohort resamples without replacement, the four primary metrics had resampling standard deviations averaging 0.010 and ranging from 0 to 0.058 across seven resampled scenarios and four methods. These summarize resampling variability, not independent biological replication. Random-label stability was summarized as mean ± s.d. across 50 partitions per cohort. Genome-wide false-positive testing comprised 180 synthetic-null scans. Nonlinear bridge comparisons used identical folds and 2,000 paired case-bootstrap resamples, with 95% confidence intervals. Simulation summaries report medians and empirical 2.5th–97.5th percentile intervals across 30 paired seeds. Feature-level inference used moderated linear models and Benjamini–Hochberg FDR control at 0.05, with effect-size thresholds and patient splits specified for each design.

Subtype labels were used for benchmark construction and evaluation, and never as correction covariates. The primary TCGA correction shared one fixed configuration across cohorts. Nested model-selection grids, simulation settings and external HK panel screening are specified in the corresponding Methods.

## Data Availability

The Cancer Genome Atlas data analysed in this study are publicly available through the National Cancer Institute Genomic Data Commons (GDC, https://portal.gdc.cancer.gov) under project identifiers TCGA-BRCA, TCGA-COAD, TCGA-GBM, TCGA-KIRC, TCGA-KIRP, TCGA-LIHC, TCGA-LUAD, TCGA-LUSC, TCGA-PRAD, TCGA-READ, TCGA-STAD, TCGA-THCA, and TCGA-UCEC. External datasets are available from the NCBI Gene Expression Omnibus (GEO) under accessions GSE4290, GSE75067, and GSE63409 [20, 38, 41].

## Code Availability

A GitHub repository containing the code is being prepared and will be published with the manuscript.

